# Activation loop phosphorylation tunes conformational dynamics underlying Pyk2 tyrosine kinase activation

**DOI:** 10.1101/2022.12.03.518986

**Authors:** TM Palhano Zanela, A Woudenberg, KG Romero Bello, ES Underbakke

## Abstract

Pyk2 is a multidomain non-receptor tyrosine kinase that undergoes a multistage activation mechanism. Activation is instigated by conformational rearrangements relieving autoinhibitory FERM domain interactions. The kinase autophosphorylates a central linker residue to recruit Src kinase. Pyk2 and Src mutually phosphorylate activation loops to confer full activation. While the mechanisms of autoinhibition are established, the conformational dynamics associated with autophosphorylation and Src recruitment remain unclear. We employ hydrogen/deuterium exchange mass spectrometry (HDX-MS) and kinase activity profiling to map the conformational dynamics associated with substrate binding and Src-mediated activation loop phosphorylation. Nucleotide engagement stabilizes the autoinhibitory interface, while phosphorylation deprotects both FERM and kinase regulatory surfaces. Phosphorylation organizes active site motifs linking catalytic loop with activation segment. Dynamics of the activation segment anchor propagate to EF/G-helices to prevent reversion of the autoinhibitory FERM interaction. We employ targeted mutagenesis to dissect how phosphorylation-induced conformational rearrangements elevate kinase activity above the basal autophosphorylation rate.

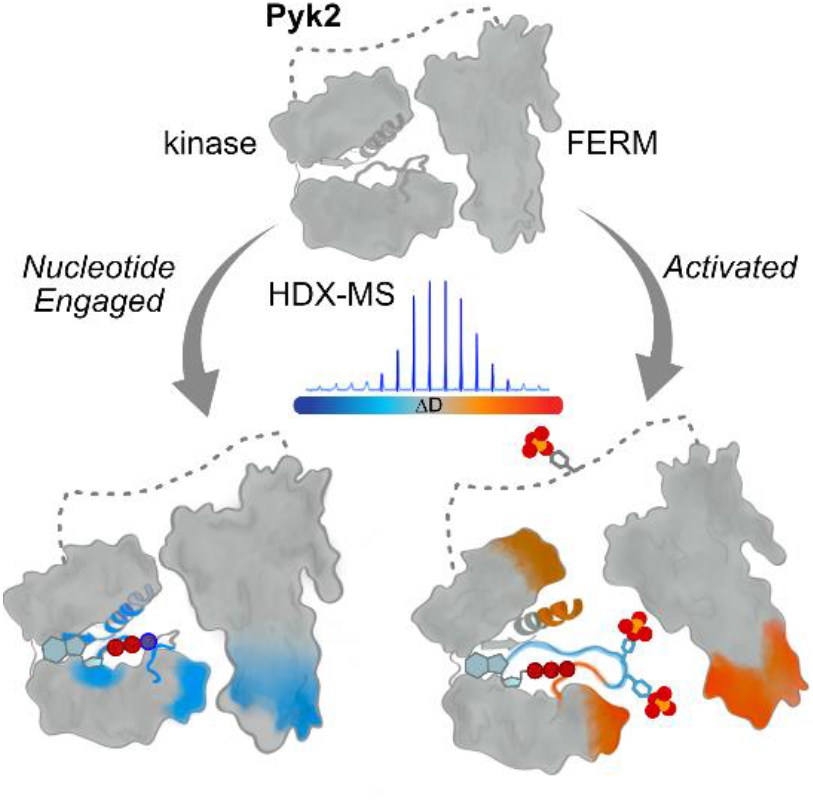

## Introduction

The signaling activities of eukaryotic protein kinases are strictly regulated by diverse, multifactor mechanisms. Kinases are maintained in a low, basal activity state by regulatory subdomains, pseudosubstrates, and proteins effectors that thwart productive catalytic conformations and substrate access. The high-activity kinase state requires strict conformational alignment for optimal phosphotransfer catalysis.^1-3^ Accessing the high-activity conformation is often dependent on phosphorylation of the activation loop, a dynamic stretch of the activation segment at the lip of the substrate docking site. The mechanisms by which regulatory features disengage to allow for activation loop phosphorylation and high-activity conformations are highly varied.^4^ Ultimately, the diversity of kinase regulatory mechanisms bestows the robustness, crosstalk, and adaptability that characterize sophisticated cellular signaling.

Active eukaryotic protein kinases generally share several interlinked conformational constraints that align catalytic residues and substrates. The conserved kinase fold is comprised of an N-lobe with a prominent C-helix and a largely alpha helical C-lobe punctuated by a catalytic loop and activation segment (Figure 1A). The N- and C-lobes of the kinase domain close to envelope an ATP substrate. The ATP nucleobase slots into and stabilizes the catalytic spine, a stack of hydrophobic residues spanning N- and C-lobes.^5^ Concomitantly, the N-lobe C-helix dips towards the active site pocket.^6^ The DFG motif at the root of the activation segment flips inward to coordinate with catalytic Mg^2+^. The shifted C-helix and DFG motif contribute sidechains to coalesce a regulatory spine linking the N-lobe and activation segment.^7^ Moreover, phosphorylated activation loop residues can ion pair with a basic cluster composed of residues contributed by the C-helix, activation segment, and catalytic loop.^8^ Thusly, activation loop phosphorylation can pin together key catalytic and substrate recognition motifs bridging N- and C-lobes. While the final catalytically productive conformations of different kinases are analogous, pathways toward activation vary considerably.

**Figure 1.**
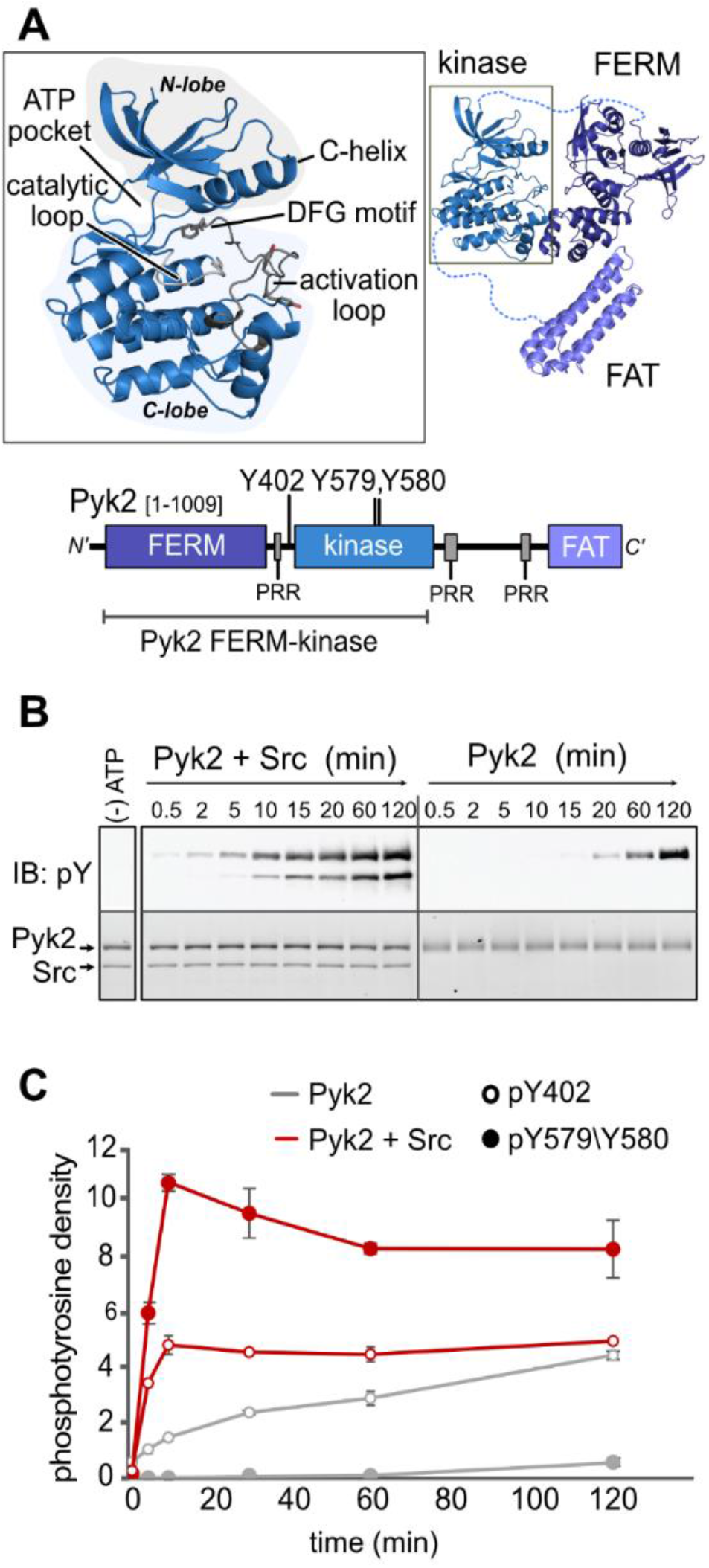
Role of Src kinase in Pyk2 phosphorylation. (A) Canonical catalytic motifs are labeled in the Pyk2 kinase domain. Higher-order Pyk2 domain organization is represented as both an AlphaFold-derived structural model and cartoon domain map. Phosphorylated tyrosine residues and proline-rich regions (PRR) are annotated. (B) Kinase activity time course of Pyk2 FERM–kinase (0.5 μM) in the presence or absence of Src (0.2 μM). Tyrosine phosphorylation was detected via Western blotting with pan-specific anti-phosphotyrosine primary antibody (PY20) (C) Site-specific phosphorylation was detected by blotting with anti-phospho-PYK2 (pY579/Y580) or anti-phospho-PTK2B (pY402). Phosphorylation levels were quantified by densitometry. Error bars represent the standard deviation of three independent reaction time courses.

Conformational dynamics are fundamental to the transition from inactive to active kinase. Time-resolved structural biology approaches such as NMR, molecular dynamics simulations, and hydrogen/deuterium exchange mass spectrometry (HDX-MS) have been invaluable for revealing conformational dynamics associated with activation.^9-14^ Here, we probe the conformational dynamics associated with activation of non-receptor tyrosine kinase proline-rich tyrosine kinase 2 (Pyk2), the lone paralog of focal adhesion kinase (FAK). We are especially interested in Pyk2 as a FAK gene duplication subject to evolutionary drift into unique regulatory responsiveness.

FAK and Pyk2 share a conserved domain organization with an N-terminal regulatory FERM (protein 4.1/ezrin/radixin/moesin) domain, a central kinase, and a C-terminal FAT (focal adhesion targeting) domain (Figure 1A). The trilobed FERM domain mediates autoinhibition by engaging with the kinase C-lobe.^15,16^ The FERM obscures access to the kinase activation loop and, presumably, blocks docking of protein substrates. Despite the autoinhibitory architectural similarities, Pyk2 diverged to respond to activational stimuli distinct from FAK. Whereas FAK is canonically activated by membrane lipid interactions and clustering at focal adhesions,^17,18^ Pyk2 has adopted Ca^2+^ sensitivity in neuronal cells.^19-21^ However, the mechanistic details of activation remain unclear, especially in Pyk2.^22^ Ultimately, activation involves autophosphorylation of a key tyrosine (FAK Y397, Pyk2 Y402) in the FERM–kinase linker. The phosphorylated linker tyrosine and a proximal proline-rich region serve as a docking site for Src kinase. Src and FAK mutually phosphorylate activation loops to achieve full activity.^23,24^ Src-mediated phosphorylation also underlies Pyk2 activation, yet the mechanistic details are ambiguous.^25,26^ The structure of the phosphorylated form of the isolated FAK kinase domain has been reported.^15^ However, no structures have been reported of the active conformation of Pyk2. Likewise, no structural models are available describing the conformational transitions of FAK or Pyk2 from autoinhibited to active, phosphorylated forms. The paucity of Pyk2 structural snapshots compelled us to explore the conformational changes driving Pyk2 activation. We reasoned that a focus on Pyk2 activation loop phosphorylation would reveal changes in kinase dynamics linking de-repression of the regulatory FERM domain with the assembly of a catalytically productive active site.

To illuminate the conformational dynamics associated with phosphorylation-induced Pyk2 activation, we integrate functional assays, HDX-MS, and targeted site-directed mutagenesis studies. We resolve the phosphorylation cascade of Pyk2 and Src by reconstituting kinase complexes and dissecting site-specific phosphorylation targets. Accordingly, we establish *in vitro* conditions for generating a fully active, phosphorylated Pyk2 FERM–kinase construct. We employ HDX-MS to map the conformational dynamics and higher-order structural changes associated with Pyk2 ATP engagement and activation loop phosphorylation. Our results reveal distinct patterns of conformational stabilization and deprotection impacting FERM and kinase regulatory interfaces, activation segment, and catalytic loop.

## Results

### Src-kinase mediates Pyk2 activation loop phosphorylation *in vitro*

We sought to resolve ambiguities regarding the role of Src in Pyk2 activation. Cell-based studies have indicated that Pyk2 autophosphorylation is either a prerequisite for Src-recruitment ^25^ or dependent on Src-mediated priming.^26^ To investigate the requirements for Pyk2 autophosphorylation, we purified the minimal regulatory Pyk2 FERM–kinase construct (residues 20-692) in a defined, unphosphorylated state (Figure 1A).^15,16^ We investigated the capacity of unphosphorylated Pyk2 FERM–kinase to autophosphorylate *in vitro*. Indeed, Pyk2 alone exhibits a basal capacity for autophosphorylation (Figure 1B). The inclusion of Src (full-length, unphosphorylated), markedly increases Pyk2 phosphorylation rates (Figure 1B). To assess whether the Src-induced increase in phosphorylation rate was associated with Pyk2 activation loop phosphorylation, we monitored site-specific phosphorylation in the Pyk2 in the presence or absence of Src. Blotting with site-specific antibodies revealed that the target of Pyk2 autophosphorylation was primarily FERM—kinase linker residue Y402, the putative Src docking site (Figure 1C and S1). Pyk2 exhibited negligible autophosphorylation of its own activation loop residues Y579 and Y580. The inclusion of Src, however, resulted in vigorous phosphorylation of Pyk2 activation loop tyrosines (Figure 1C and S1). Our results confirm that unphosphorylated Pyk2 exhibits basal autophosphorylation activity, independent of Src activity. Pyk2 activation loop (Y579/Y580) phosphorylation, however, is Src-induced, consistent with cell-based studies.^25^ Furthermore, phosphorylated, high-activity Pyk2 promotes reciprocal phosphorylation of Src (Figure 1B).

### Nucleotide-binding orders regulatory elements in the Pyk2 kinase domain

Our investigations into the kinase activity requirements of Pyk2 revealed that Src can drive the FERM–kinase construct to the fully phosphorylated, high-activity form *in vitro*. With access to both autoinhibited (unphosphorylated) and high-activity (fully phosphorylated) Pyk2 states, we sought to define the conformational dynamics associated with stages of Pyk2 activation. We began by mapping the conformational changes associated with nucleotide substrate binding in the autoinhibited conformation. We employed HDX-MS to assess nucleotide-induced structural perturbations by comparing H/D exchange rates of unbound Pyk2 FERM–kinase with Pyk2 bound to a non-hydrolyzable ATP analog (AMPPNP). HDX-MS reports on protein dynamics by monitoring the rates of protein backbone amide proton exchange with deuterated solvent. H/D exchange rates are exquisitely sensitive to secondary structure, hydrogen bonding, conformational dynamics, and solvent exposure. As such, exchange rate perturbations are useful for mapping of local protein dynamics, interfaces, and allostery.^27^

AMPPNP-binding induced localized patterns of exchange rate suppression in key catalytic and regulatory regions of Pyk2 FERM–kinase (Figure 2 and S2). The kinase ATP-binding pocket exhibited significant exchange rate decreases in both the glycine-rich loop and the αC-helix. The glycine-rich loop is a conserved kinase motif that interacts with ATP substrate β- and γ-phosphates via backbone amide interactions. The αC-helix includes a key catalytic residue, a glutamate (Pyk2 E474) that ion pairs with an N-lobe β-sheet lysine (Pyk2 K457) to promote catalytically productive alignment of the ATP α- and β-phosphates. AMPPNP-binding also induced exchange rate decreases in the N- and C-terminal peripheries of the activation segment, including the DFG and APE motifs. The DFG motif is a conformationally switchable sequence that includes the aspartate responsible for coordinating catalytic Mg^2+^ and the phenylalanine that integrates into the regulatory spine.^28^ The APE motif (in Pyk2, SPE) roots the activation segment and may participate in protein substrate docking.

**Figure 2.**
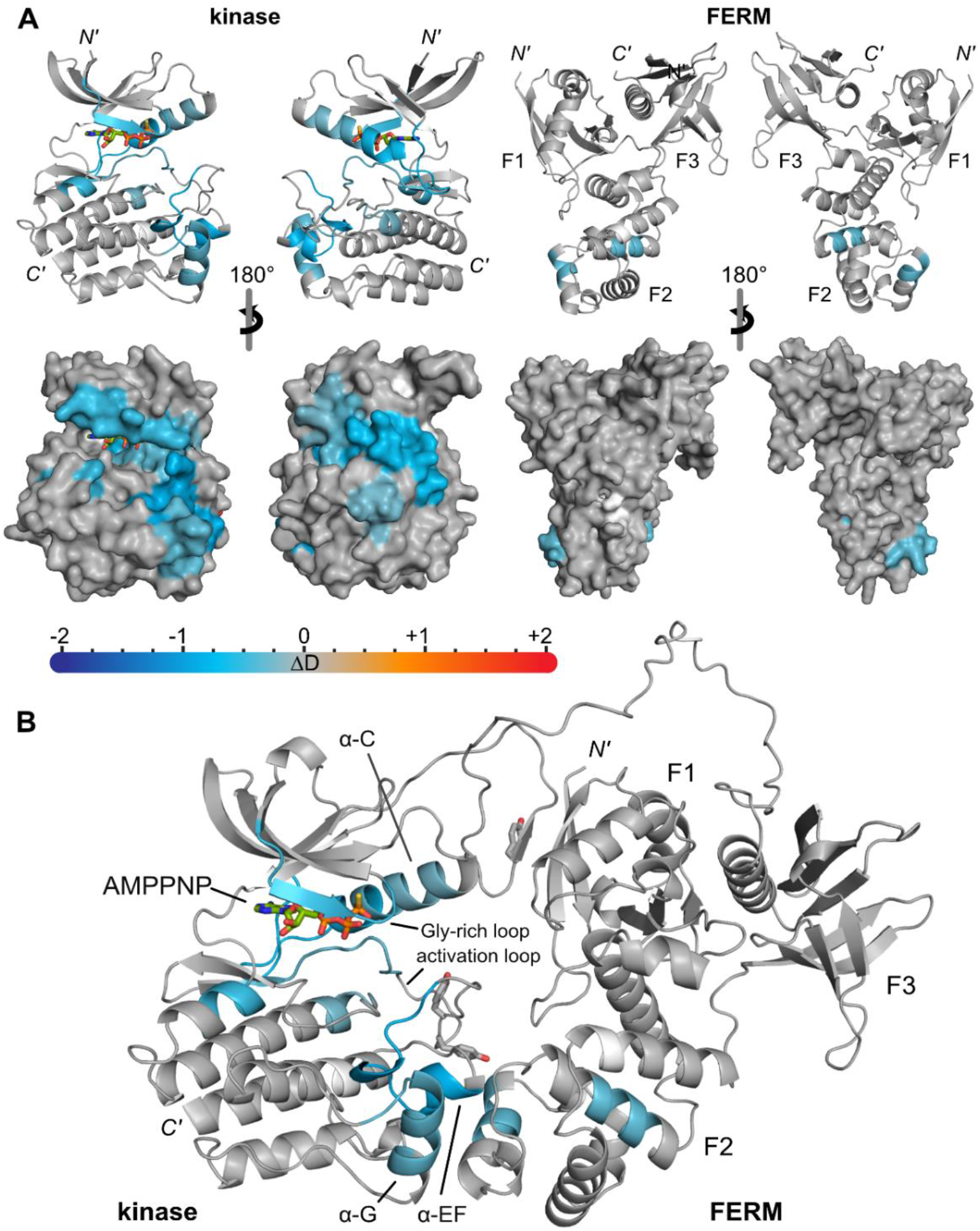
HDX-MS reveals nucleotide-induced stabilizations in Pyk2 FERM–kinase. (A) Exchange rate perturbations of AMPPNP-bound Pyk2 FERM–kinase relative to the unbound state are mapped to AlphaFold models of isolated Pyk2 FERM and kinase domains (AlphaFold model AF-Q14289-F1).^29^ N′- and C′-termini and FERM subdomains (F1−F3) are annotated. Exchange rate perturbations are reported as the average difference in deuterium incorporation (ΔD) at time points approximating the midpoint of exchange. Regions exhibiting significant (p < 0.005) exchange rate perturbations are color-coded according to the scale bar. Significance was assessed with a two-tailed, unpaired Student’s *t* test. Slower exchange in the AMPPNP-bound Pyk2 FERM–kinase relative to apo are mapped in blue hues, while faster exchange is mapped in red hues. Non-significant differences are colored gray. (B) HDX-MS exchange rate perturbations mapped to the structural model of the autoinhibited Pyk2 FERM–kinase (AF-Q14289-F1).

Interestingly, AMPPNP-binding also impacted regulatory surfaces distant from the ATP-binding pocket. The kinase αEF and αG helices exhibited AMPPNP-induced exchange rate suppression. In addition, the F2 subdomain of the autoinhibitory FERM domain displayed patches of decreased exchange. The kinase αEF and αG helices and FERM F2 subdomain are the primary interaction surfaces stabilizing the autoinhibited conformation of FAK and Pyk2 FERM–kinase (Figure 2B).^15,16^

### Src-induced Pyk2 phosphorylation perturbs the FERM–kinase interface and stabilizes kinase catalytic motifs

Next, we assessed the conformational dynamics associated with high-activity, fully phosphorylated Pyk2. We leveraged conditions optimized for reciprocal Pyk2 and Src phosphorylation (Figure 1C) to generate Pyk2 FERM–kinase with phosphorylated activation loop and FERM–kinase linker. The fully phosphorylated Pyk2 construct was saturated with excess ATP for comparison with the AMPPNP-bound, unphosphorylated Pyk2 FERM–kinase. HDX-MS time courses revealed significant patterns of exchange rate perturbation induced by Pyk2 phosphorylation (Figure 3A and S3). Pyk2 phosphorylation induced deprotection (i.e., increased exchange rates) in the FERM domain, while discrete regions of protection and deprotection were evident in the kinase domain.

**Figure 3.**
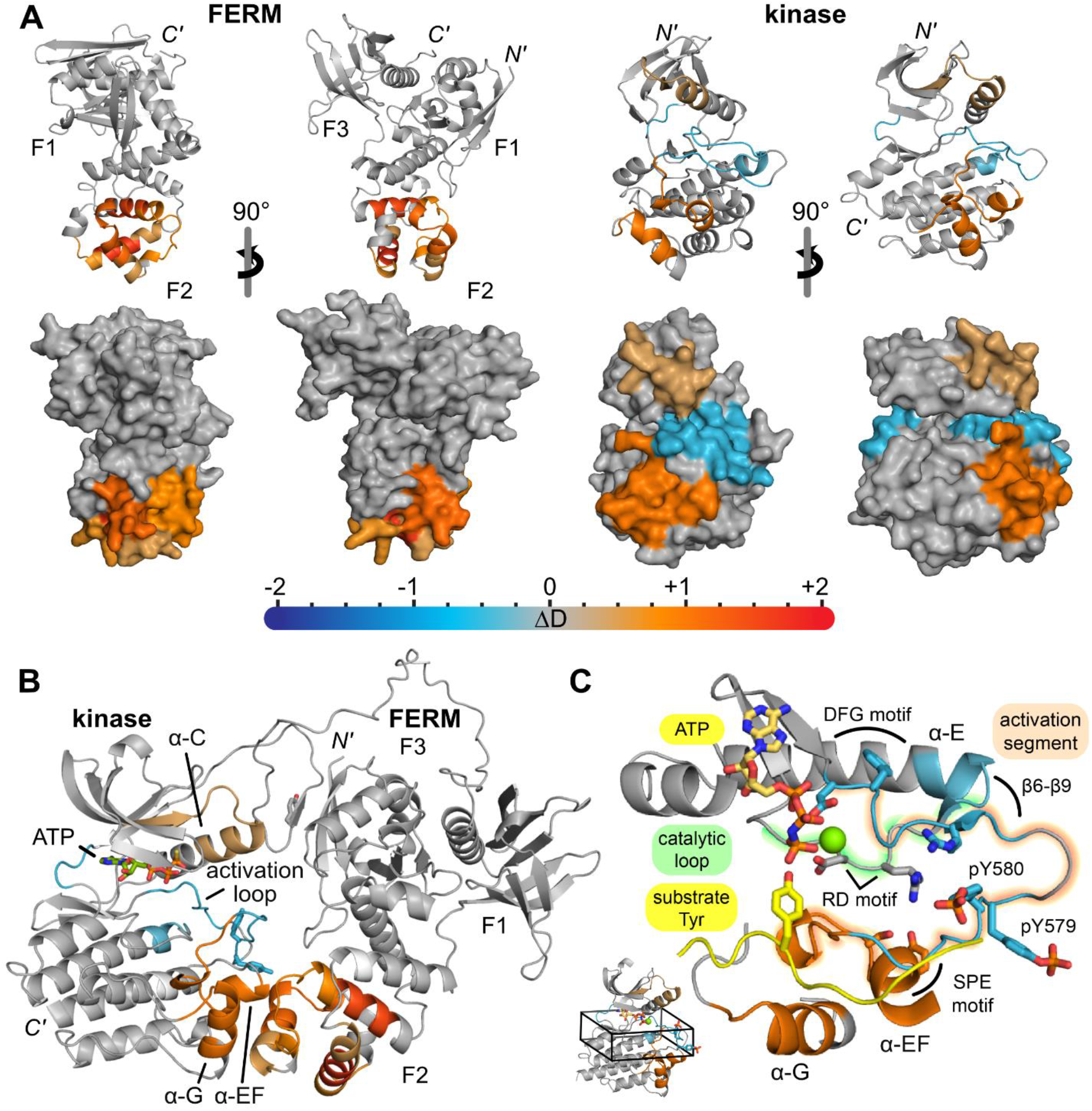
HDX-MS illuminates the conformational dynamics of high-activity, fully phosphorylated Pyk2 FERM–kinase. (A) Exchange rate perturbations of ATP-bound, phosphorylated Pyk2 FERM–kinase compared to the unphosphorylated, AMPPNP-bound state are mapped to AlphaFold models of isolated Pyk2 FERM and kinase domains (AlphaFold model AF-Q14289-F1).^29^ N′- and C′-termini, FERM subdomains (F1−F3) are annotated. Exchange rate perturbations are reported as the average difference in deuterium incorporation (ΔD) at time points approximating the midpoint of exchange. Regions exhibiting significant (p < 0.005) exchange rate perturbations are color-coded according to the scale bar. Significance was assessed with a two-tailed, unpaired Student’s *t* test. Slower exchange in the phosphorylated, ATP-bound Pyk2 FERM–kinase relative to the AMPPNP-bound form is reported in blues, while faster exchange is shown in reds. Non-significant differences are colored gray. (B) HDX-MS exchange rate perturbations mapped to the structural model of the autoinhibited Pyk2 FERM–kinase (AlphaFold model AF-Q14289-F1). (C) Zoomed view of the Pyk2 kinase active site region with mapped exchange rate perturbations. The active conformation of Pyk2 with docked protein tyrosine substrate was predicted with AlphaFold2 using ColabFold (ver. 1.0).^30^ ATP substrate and phosphotyrosines (pY579 and pY580) are modeled based on FAK kinase (PDB entry 2j0l). Additional annotations include beta strands β6 and β9, the conserved RD motif, and the DFG and SPE motifs denoting the N- and C-terminal ends of the activation segment.

The most prominent deprotection induced by Pyk2 phosphorylation localizes to the autoinhibitory interface comprising the F2 FERM subdomain and kinase C-lobe αEF and αG helices (Figure 3B). The striking deprotection at the autoinhibitory FERM and kinase surfaces is a reversal of the apparent stabilization of the autoinhibitory interface induced by nucleotide binding (Figure 2B). The increased conformational dynamics induced by phosphorylation throughout the Pyk2 autoinhibitory interface are consistent with the apparent incompatibility of activation loop phosphorylation and FERM engagement observed in FAK.^15^

Phosphorylation induced further changes to conformational dynamics throughout the kinase domain. The N-lobe αC-helix and β3 strand exhibited modest increases in exchange rate (Figure 3A), a reversal of the nucleotide-binding effect (Figure 2A). Activation segment conformational dynamics were also impacted by phosphorylation. The region around the DFG motif was further stabilized beyond the degree attributed to nucleotide binding. To estimate dynamics at the site of activation loop phosphorylation, we compared exchange rates of the peptide including phosphorylated Y579/Y580 with the same unphosphorylated peptide from the unphosphorylated, AMPPNP-bound sample. Activation loop phosphorylation apparently decreased exchange locally around Y579/Y580. However, the neighboring sequence at the C-terminal root of the activation segment– including the SPE motif– exhibited sharply increased exchange rates. Phosphorylation also induced lower exchange at the N-terminal end of the catalytic loop, including the tip of the preceding αE-helix.

### FERM–kinase Pyk2 interface is stabilized by hydrophobic and polar interactions

Our HDX-MS analysis revealed that the Pyk2 autoinhibitory interface between the FERM F2 and kinase C-lobe is stabilized by nucleotide binding (Figure 2B) and strikingly deprotected upon activation loop phosphorylation (Figure 3B). We sought to refine the interface mapped by HDX-MS by testing the regulatory contribution of specific residues. FAK and Pyk2 both exhibit hydrophobic residues at the core of the interface critical to stabilizing the autoinhibitory conformation.^15,16^ Both kinase domains project an αEF-helix phenylalanine (F599 in Pyk2) that nestles into a hydrophobic pocket in the FERM F2 subdomain. Exchange rate perturbations due to Pyk2 phosphorylation expand beyond this hydrophobic interaction “hot spot” to a network of peripheral charged residues, including D189, K197, and E207 in the FERM and R600, R601, and E639 in the kinase (Figure 4A). Notably, HDX-MS alone cannot distinguish conformational dynamics due to direct interaction from indirect allosteric perturbations. We aimed to test whether the apparent peripheral electrostatic ring directly contributed to the regulatory FERM— kinase interface. We reasoned that if electrostatics stabilized the autoinhibited conformation, Pyk2 activity could be de-repressed with increasing ionic strength. Indeed, Pyk2 FERM–kinase autophosphorylation increases with increasing (0 – 300 mM NaCl) ionic strength (Figure S4).

**Figure 4.**
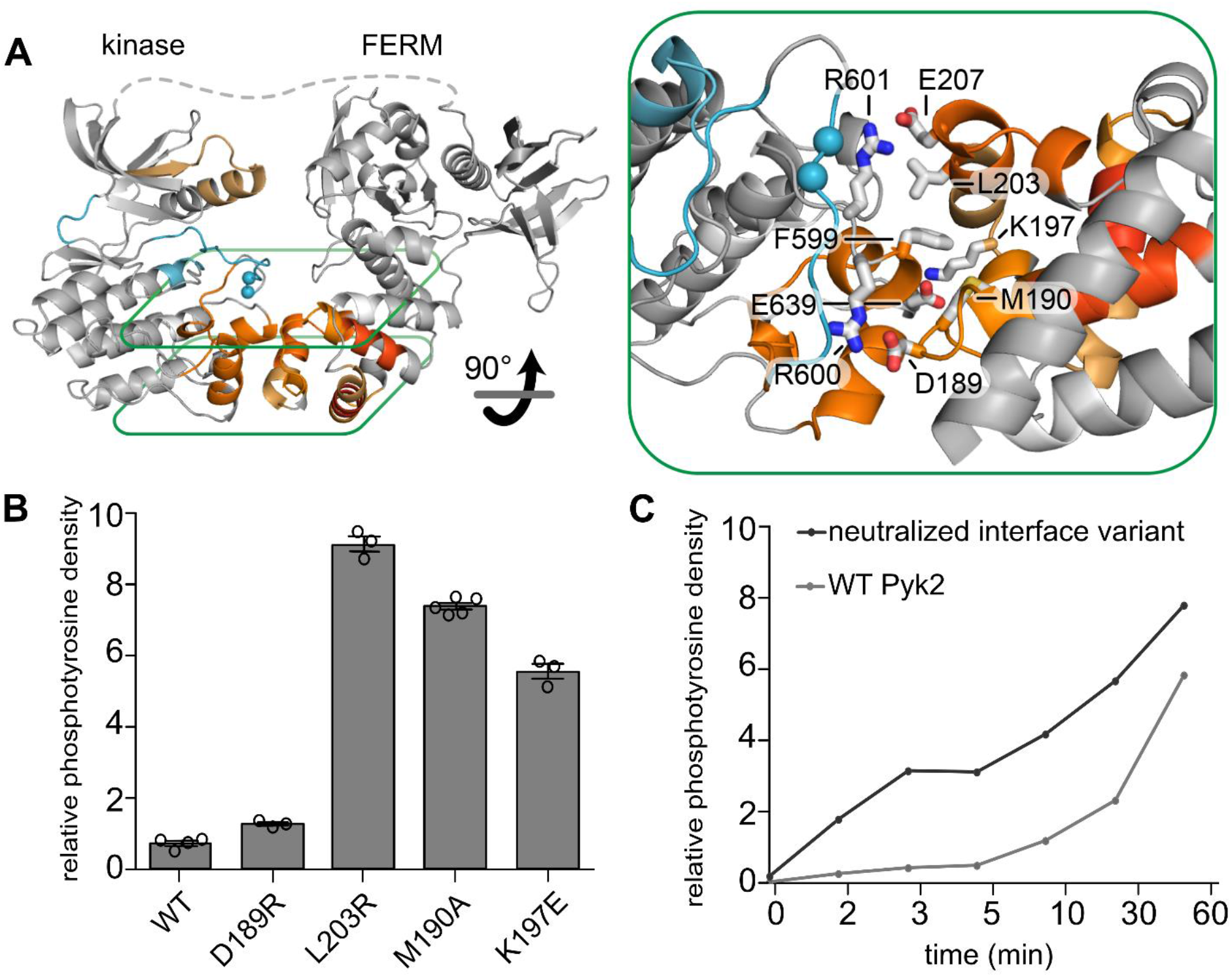
The FERM—kinase interface is stabilized by a hydrophobic core and surrounding electrostatic ring. (A) Phosphorylation-induced deprotection of the FERM–kinase interface is highlighted and color-coded as in Figure 3. Exchange rate increases map to core hydrophobic residues and a periphery of charged residues across kinase and FERM domains (inset). (B) Charge inversion and sidechain-pruned interface variants impact Pyk2 FERM–kinase autophosphorylation. Autophosphorylation (5 min time point) was assessed by immunoblotting with site-specific phospho-Y402 primary antibody. Activity assay replicates (n = 3–5) represent independent reactions. Error bars signify standard deviation. (C) Neutralizing interface charges via site-directed mutagenesis (D189S, K197S, E207S, R600S, R601S, E639S) de-represses Pyk2 FERM–kinase autophosphorylation, as measured by immunoblotting with site-specific phospho-Y402 antibody.

We further explored residue-specific contributions by designing and testing putatively disruptive variants (D189R, M190A, K197E, and L203R) at the periphery of the interface mapped by HDX-MS. We found that most of the Pyk2 variants exhibited significantly increased autophosphorylation activity compared with the WT protein *in vitro* (Figure 4B). We also tested how neutralization of the electrostatic ring impacted Pyk2 autoinhibition by generating a Pyk2 FERM–kinase construct with serines substituting for the peripheral charged residues FERM residues (D189S, K197S, E207S) and the opposing kinase residues (R600S, R601S, and E639S). The autophosphorylation activity of the Pyk2 neutralized interface variant is strikingly higher than the WT Pyk2 FERM–kinase (Figure 4C). Collectively, the HDX mapping and interface variants indicate that the autoinhibitory surface is stabilized by peripheral electrostatic residues surrounding the hydrophobic interaction core. Furthermore, this expansive FERM—kinase interface is apparently overcome and largely deprotected upon activation loop phosphorylation.

### Src-mediated activation loop phosphorylation stabilizes key Pyk2 active site motifs

Phosphorylated Pyk2 FERM–kinase exhibits significant decreases in conformational dynamics at highly conserved motifs of the activation segment and catalytic pocket (Figure 3C). Notably, a peptide sampling the phosphorylated residues Y579 and Y580 exhibited lower exchange relative to the unphosphorylated treatment, suggesting conformational constraint. The apparent decrease in conformational dynamics around the phosphotyrosines was accompanied by decreased exchange at the N-terminal anchor of the activation segment and sequence adjoining the catalytic loop (Figure 5A). These regions include arginine residues (R548 and R572 in Pyk2) that serve as an ion-pairing “basic pocket” for phosphorylated activation loop residues in many kinases.^8^ Indeed, a phosphotyrosine pairs with the corresponding FAK arginines in the reported structure of a phosphorylated FAK kinase truncation.^15^ We hypothesized that the phosphorylation-induced exchange rate suppressions of the N-terminal activation segment and catalytic loop are associated with conformational constraints due to ion pairing between basic pocket residues and phosphotyrosine.

**Figure 5.**
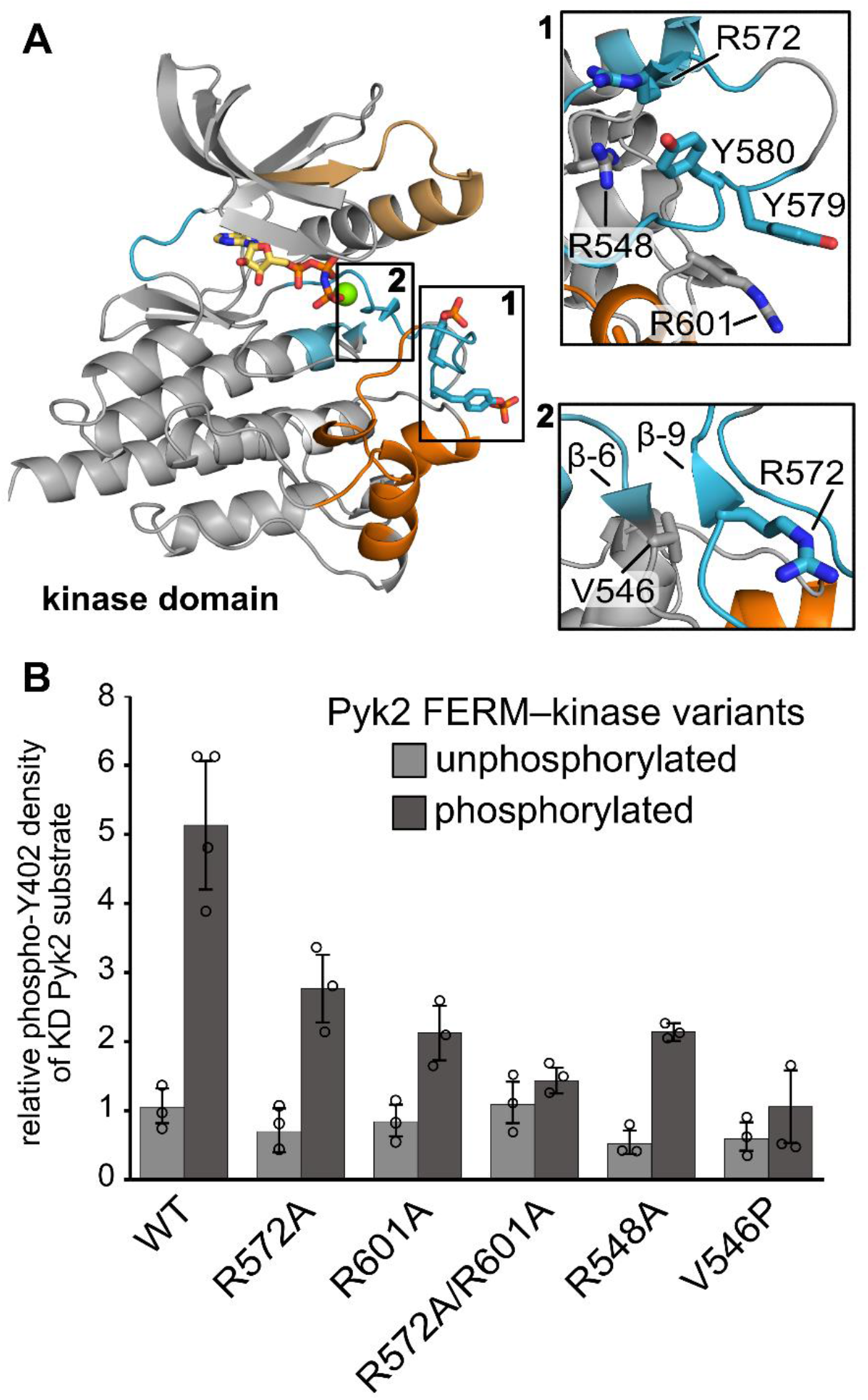
Phosphotyrosine ion pairing and beta sheet formation stabilize the activation conformation of Pyk2. (A) Phosphorylation-induced exchange rate perturbations of the kinase domain are color-coded as in Figure 3. The active conformation of Pyk2 is modeled with AlphaFold2 using ColabFold.^30^ ATP substrate and phosphotyrosines (pY579 and pY580) are modeled based on FAK kinase (PDB entry 2j0l). Insets highlight Pyk2 arginines proximal to the activation loop tyrosines (inset 1), and beta strands β6 and β9 (insert 2). (B) Kinase activity of unphosphorylated and fully phosphorylated Pyk2 FERM–kinase variants (0.02 µM) was assessed by monitoring Y402 phosphorylation (45 min) of a kinase-dead (KD) Pyk2 construct (1 µM). Phosphotyrosine density was quantified by densitometry after Western blotting with primary antibody specific to phospho-Y402. Activity assay replicates (n = 3–4) represent independent reactions. Error bars signify standard deviation.

We investigated whether the basic pocket arginines contribute electrostatic interactions necessary for the high-activity kinase conformation upon activation loop phosphorylation. To systematically remove basic pocket charges, we engineered alanine variants at three key residues, R572, R548, and R601 (Figure 5A). R572A is located within the activation segment anchor exhibiting phosphorylation-induced exchange rate suppression. The R548A variant ablates the HRD motif arginine adjacent to the catalytic loop region ordered by phosphorylation. We also generated R601A and R572A/R601A to test a possible electrostatic interaction with the second activation loop phosphotyrosine. To compare activities of variants with defined phosphorylation states, we monitored Y402 phosphorylation of a kinase dead Pyk2 FERM-kinase construct added in 50-fold excess (Figure S5). All four arginine variants preserved basal phosphorylation activity, comparable to the wild-type Pyk2 FERM–kinase construct (Figure 5B). We next tested whether arginine substitutions impacted kinase activation by phosphorylating the activation loop of each variant with Src. Phosphorylation of R548A, R572A, and R601A significantly increased kinase activity over the basal level. However, each phosphorylated arginine variant failed to reach full kinase activation, exhibiting half the activity of the phosphorylated wild-type construct (Figure 5B). Furthermore, phosphorylation of the double variant R572A/R601A induced negligible activation over basal activity. Ultimately, all three arginine residues in the vicinity of phosphorylation-induced exchange rate suppressions were important to stabilizing the high-activity conformation upon activation loop phosphorylation.

In addition to contributing basic pocket arginines, the regions near the DFG motif and catalytic loop comprise the β6 and β9 strands that zipper into a short, antiparallel β-sheet, a hallmark of active kinases.^8^ The β6 and β9 strands exhibited phosphorylation-induced protection, consistent with secondary structure formation (Figure 5A). Notably, the corresponding sequences in the autoinhibited FAK FERM–kinase are splayed apart.^15^ To investigate the correlation between the putative formation of the β6–β9 sheet and the high-activity conformation, we designed a Pyk2 variant (V546P) to disrupt the hydrogen bonding between strands (Figure 5A). Introduction of the proline within β6 did not significantly impair basal kinase activity (Figure 5B). However, Src-mediated activation loop phosphorylation of the V546P variant failed to elevate kinase activity above the basal rate (Figure 5B). These observations indicate that formation of the stabilizing beta sheet linking activation segment to catalytic loop is a strict prerequisite for full Pyk2 activation.

## Discussion

FAK and Pyk2 undergo a complex, multistage activation process involving large conformational rearrangements, Src recruitment, and multiple phosphorylations. Dissecting the conformational dynamics underlying the maturation of the kinase activation complex is important for understanding both shared and divergent regulatory mechanisms in FAK and Pyk2. Here, we reveal the mechanisms and conformational dynamics associated with Src-mediated activation of Pyk2. By integrating HDX-MS and functional assays with structural models of Pyk2 domains, we illuminate allosteric linkages between phosphorylated activation loop, catalytic core, and regulatory interfaces.

Reconstitution of the Pyk2/Src activation complex *in vitro* revealed that Pyk2 exhibits an intrinsic capacity to autophosphorylate linker residue Y402, independent of activation loop phosphorylation (Figure 1). Reciprocal phosphorylation of Pyk2 and Src activation loops confers maximal kinase activity. Notably, Pyk2 activation can be recapitulated *in vitro* without clustering or Ca^2+^-triggering,^20,21^ indicating that the autoinhibitory FERM interaction is dynamic. However, *in vitro* reconstitutions exclude endogenous cellular phosphatases that would otherwise suppress basal autophosphorylation.^31^ Indeed, the recently reported Ca^2+^/calmodulin-induced dimerization of Pyk2 via the intrinsically disordered kinase—FAT linker is likely a critical factor in outcompeting the activity of cellular phosphatases for robust activation *in vivo*.^19^

We capitalized on the reconstitution of Pyk2 with defined phosphorylation states by exploring the conformational dynamics associated with nucleotide substrate binding and activation loop phosphorylation. Reported high-resolution structures of Pyk2 and FAK kinase domains are invaluable resources, but static snapshots may obscure the dynamics responsible for activation. For example, the structures of autoinhibited FAK FERM–kinase and phosphorylated FAK kinase both exhibit signatures of active kinase conformations (RMSD ∼0.8 Å), including an active site-engaged αC-helix and assembled regulatory spine.^15^ In contrast, a structure reported of truncated Pyk2 kinase bound to ATPγS has a broken regulatory spine with the DFG phenylalanine slipped into the active site pocket.^32^ Using HDX-MS, we introduced a temporal dimension to the structural models to map activation stages of Pyk2 FERM–kinase.

HDX-MS revealed that nucleotide substrate binding stabilizes the glycine-rich loop, αC-helix, and the activation segment of Pyk2 FERM–kinase (Figure 2). Notably, autoinhibitory engagement of the FERM domain does not preclude nucleotide binding. While N- and C-lobes are closed in both crystallography-derived structures of autoinhibited and active FAK kinase,^15^ exchange protection across both lobes of Pyk2 kinase indicates that nucleotide binding decreases inter-lobe conformational dynamics. Interestingly, HDX-MS analysis of nucleotide binding in ERK2 kinase suggests contrasts with Pyk2.^33^ Nucleotide binding only impacts the N-lobe in inactive, unphosphorylated ERK2, while engagement of both N- and C-lobes requires the active, phosphorylated state. Strikingly, Pyk2 nucleotide binding also stabilizes the autoinhibitory interface in both FERM and kinase domains. FRET-based conformational sensors in FAK also suggest that ATP-binding biases FAK FERM–kinase toward a compacted, autoinhibitory conformation.^34^

HDX-MS and targeted mutagenesis also revealed key Pyk2 conformational changes resulting from activation loop phosphorylation (Figure 3). Conspicuous deprotection of the autoinhibitory interface indicates disengagement of the FERM domain, mirroring the incompatibility of FERM docking with activation loop phosphorylation observed in FAK.^15^ Interestingly, dual phosphorylation of the activation loop is sufficient to outcompete the extensive 650 Å^2^ FERM— kinase interface composed of interdigitating hydrophobic residues and a periphery of electrostatic interactions (Figure 4). The concomitant increase in dynamics of both the activation segment C-terminal anchor and αEF/G-helices suggests allosteric linkage between the phosphorylated activation loop and autoinhibitory FERM interface. Similarly, TEC family kinase regulatory interactions between PHTH domain and kinase converge on the G-helix region and allosterically communicate with the active site cleft.^35^

In addition to dislodging the FERM domain, activation loop phosphorylation stabilizes key active site motifs. Catalytic competency appears to arise from subtle conformational constraints, as large-scale conformational rearrangements are not obvious in comparisons of autoinhibited vs. active FAK kinase structures.^15^ Exchange rate perturbations implicated phosphotyrosine interactions with a cluster of basic residues as well as formation of a putative beta sheet (β6/β9) linking of catalytic loop and activation segment (Figure 3C). Indeed, selectively perturbing these regions with point mutations ablates the phosphorylation-induced high activity state without impairing the basal autophosphorylation activity (Figure 5). Nevertheless, intrinsic exchange rates of the activation segment and N/C-lobe hinge remain high, suggesting that the active conformation retains high intrinsic dynamics. Kinase inter-lobe dynamics are likely critical for the catalytic cycle and substrate/product exchange.^8,36^

Ultimately, activation loop phosphorylation spurs increased conformational dynamics in the Pyk2 kinase C-lobe (e.g., αEF- and αG-helices) attributable to FERM dissociation, while pre-organizing the catalytic pocket. Nucleotide substrate binding stabilizes a subset of active site regions, whereas activation loop phosphorylation further constrains catalytic motifs. Indeed, time-resolved, solution-based structural approaches have been critical in highlighting the important shifts in conformational landscapes organizing the active sites of diverse kinases.^10,11,13,36^

## Methods

### Plasmid Constructs

The Pyk2 gene was synthesized from gBlocks (Integrated DNA Technologies) with codon usage optimized for recombinant expression in *E. coli*. Pyk2 FERM–kinase (residues 20-692) was subcloned in-frame with an N-terminal H_6_-SUMO affinity/solubility tag in expression vector pET-H6-SUMO-TEV-LIC (1S), a gift from S. Gradia (Addgene, 29659). The resulting H6-SUMO-FERM–kinase under T7 promoter control was designated pNAM004. Pyk2 FERM– kinase mutants in pNAM004 were generated using the QuikChange (Agilent Genomics) site-directed mutagenesis strategy. The YopH tyrosine phosphatase expression vector pESU009 was generated by cloning the YopH phosphatase domain (residues 177-468) into pACYCDuet-1 (Novagen) under transcriptional control of an attenuated *tac* promoter.^37^ All constructs were confirmed by DNA sequencing.

### Protein Purification

#### Pyk2 FERM–kinase purification

pNAM004 and mutants were co-transformed into NiCO21(DE3) *E. coli* cells with pESU009 encoding phosphatase YopH to ensure homogenous, dephosphorylated product. Purification of Pyk2 FERM–kinase and its variants was performed as previously described,^16^ with the following modifications. After protein expression and harvesting, cells were lysed by sonication in 50 mM Tris pH 8, 25 mM NaCl, 20 mM imidazole, 1 mM phenylmethylsulfonyl fluoride (PMSF), 5 mM EDTA, 5% glycerol, 2.5 mM β-mercaptoethanol (βME) supplemented with protease inhibitor cocktail II (Research Products International). Lysate was cleared of cell debris by ultracentrifugation at 80,000 ×g (Beckman MLA-50 rotor) for 15 min at 4 °C. Pyk2 constructs were enriched on Ni-NTA affinity resin (HisTrap HP 5 mL, Cytiva), and the H_6_-SUMO tag was removed via H_6_-Ulp1 protease treatment and a subtractive passage through Ni-NTA. The resulting Pyk2 constructs were subsequently concentrated using a 10k molecular weight cutoff (MWCO) PES Spin-X centrifugal concentrator (Corning) and further purified by gel filtration chromatography (GFC) using a Superdex 200 10/300 (GE Healthcare) column pre-equilibrated in GFC buffer [50 mM HEPES pH 7.4, 150 mM NaCl, 5% glycerol, 5 mM TCEP]. The purified proteins were aliquoted, snap frozen in liquid N_2_, and stored at -80 °C.

#### Src purification

H_6_-SUMO-Src (residues 1-536) was co-expressed with YopH in NiCO21(DE3) cells for 16 hours, shaking at 18 °C. Cells were harvested by centrifugation and stored at 18 °C until purification. Cell pellets were thawed on ice in Src Binding Buffer [50 mM Tris pH 8, 500 mM NaCl, 25 mM imidazole, 5% glycerol, 2.5 mM βME] supplemented with 2 mM EDTA and 1 mM PMSF. All subsequent purification steps were performed at 4 °C. Cells were lysed by sonication, and lysate was cleared by centrifugation at 20,000 ×g for 30 min.

Cleared lysate was supplemented with 5 mM MgCl_2_. H_6_-SUMO-tagged Src was enriched on Ni-NTA (5 mL HisTrap, Cytiva) pre-equilibrated in Src Binding Buffer. The Ni-NTA column was washed with five column volumes of Src Binding Buffer followed by elution with 50 mM Tris pH 8, 500 mM NaCl, 250 mM imidazole, 5% glycerol, 2.5 mM βME. Elution fractions containing H_6_-SUMO-Src were pooled and treated with H_6_-Ulp1 for proteolytic H_6_-SUMO tag cleavage while dialyzing into 50 mM Tris pH 8, 100 mM NaCl, 5% glycerol, 2.5 mM DTT. Protease and tag were removed with a subtractive passage through the Ni-NTA column. Column flow-through was concentrated via centrifugal concentrator (10k MWCO) and applied to a Superdex 200 10/300 (GE Healthcare) gel filtration column pre-equilibrated in 50 mM HEPES pH 7.4, 100 mM KCl, 5% glycerol, 0.8 mM TCEP. Purified Src was concentrated and snap frozen in liquid N_2_ for storage at -80 °C.

#### Kinase Assays

Kinase reactions were performed using a final concentration of 0.5 μM for WT or variant Pyk2 FERM–kinase variants in a kinase buffer consisting of 50 mM HEPES pH 7.4, 150 mM NaCl, 8 mM MgCl_2_, 5% glycerol. Kinase reactions were initiated by addition of 4 mM ATP (final concentration) and quenched at various time points with the addition of 10 mM EDTA (final concentration) followed by heat denaturation at 90 °C for 60 sec. Phosphotyrosine production was assessed via anti-phosphotyrosine Western or dot blotting using primary mouse anti-phosphotyrosine antibody (PY20, Thermo Scientific) followed by goat anti-mouse HRP conjugated secondary antibody (GOXMO HRP, Novex). Site-specific anti-phosphotyrosine blotting was performed with anti-phospho-PTK2B pTyr402 (SAB4300173, Sigma) or anti-phospho-PYK2 pTyr579/pTyr580 (44-636G, Invitrogen) primary antibodies with the goat anti-rabbit HRP-conjugated secondary antibody (GOXRB HRP, Novex). Blots were imaged with an enhanced chemiluminescent substrate (Pierce). Phosphotyrosine density was quantified by densitometry (Image Lab, Bio-Rad).

Kinase activity of unphosphorylated or fully phosphorylated WT Pyk2 FERM–kinase or variants (R548A, R572A, R501A, R572A/R601A, and V546P) was assessed by monitoring phosphorylation of a target kinase-dead (K475A) Pyk2 construct (residues 20-729/868-877) engineered to resolve distinctly from FERM–kinase via SDS-PAGE. Fully phosphorylated Pyk2 FERM–kinase was obtained by Src-mediated phosphorylation of Y579/Y580. WT Pyk2 FERM– kinase or variants (0.5 µM) were incubated with His_6_-SUMO-Src (0.15 µM) and 4 mM ATP in kinase buffer for 90 min. His_6_-SUMO-Src was subsequently removed from the reaction mixture via two cycles of depletion by incubation with HisPur Ni-NTA Superflow agarose affinity resin (Thermo Scientific). Complete removal of His_6_-SUMO-Src was validated by Western blotting.Kinase assays samples consisted of unphosphorylated or fully phosphorylated Pyk2 FERM– kinase variants (0.02 µM) incubated with a fifty-fold excess of kinase-dead Pyk2 (1 µM) in kinase buffer. Kinase reactions were initiated by adding 4 mM ATP (final concentration) and quenched with SDS-PAGE loading dye and 10 mM EDTA (final concentration) after 45 min. Phosphorylation of kinase-dead Pyk2 was assessed via anti-phosphotyrosine Western blotting using anti-phospho-PTK2B pTyr402 antibody (SAB4300173, Sigma).

#### HDX-MS Mapping

HDX time-courses were performed in parallel for the following three treatments: Src/ATP-treated, phosphorylated Pyk2 FERM–kinase; AMPPNP-bound Pyk2 FERM–kinase; and untreated, autoinhibited Pyk2 FERM–kinase. Phosphorylated Pyk2 FERM–kinase was generated by pre-treatment with sub-stoichiometric Src (3:1 Pyk2:Src) and 4 mM ATP. AMPPNP-bound Pyk2 was pre-incubated with 4 mM AMPPNP (Sigma). Autoinhibited Pyk2 was incubated with buffer. All pre-exchange treatments were performed for 30 min at 22 °C in 50 mM HEPES pH 7.4, 150 mM NaCl, 5% glycerol, 5 mM TCEP, 8 mM MgCl_2_ (final concentrations).

Deuterium exchange reactions (65 uL) were initiated with a ten-fold dilution of 20 pmol of Pyk2 FERM–kinase from each pretreatment into D_2_O exchange buffer composed of 50 mM HEPES pD 7.4, 100 mM NaCl, 5mM MgCl_2_, 5mM DTT and 90% D_2_O (final concentrations). Exchange reaction mixtures were incubated at 22 °C. At various time points (12 s, 1 min, 2.5 min, 11 min, and 122 min), labeling reactions were quenched to pH 2.5 by addition of 20% TFA, 0.15% n-dodecylphosphocholine (fos-12; Avanti Polar Lipids) and snap-frozen in liquid N_2_. Quenched exchange reaction mixtures were stored at −80 °C until LC-MS analysis. All exchange reactions were performed as three technical replicates. For HDX-MS analysis, frozen exchange samples were quickly thawed and immediately injected (50 μL) into a temperature-controlled (4 °C) ACQUITY UPLC M-class HDX platform coupled in-line to an ESI-Q-TOF Synapt G2-Si instrument (Waters). Mobile phases consisted of solvents A (HPLC-grade aqueous 0.1% formic acid) and B (HPLC-grade acetonitrile and 0.1% formic acid). Samples were digested into uniquely identifiable peptide fragments by flow through an in-line Enzymate BEH-immobilized pepsin column (5 μm particle, 300 Å pore, 2.1 × 30 mm; Waters) at 25 °C with a flow rate of 50 μL/min with 100% solvent A. Peptide products accumulated on an Acquity UPLC BEH C18 VanGuard trap column (1.7 μm, 130 Å, 2.1 mm × 5 mm; Waters) and desalted with 100% solvent A for 0.3 min at a flow rate of 120 μL/min. Pepsin-derived peptides were subsequently resolved on an Acquity UPLC BEH C18 analytical column (1.7 μm, 130 Å, 1 × 100 mm; Waters) using a 7 min linear gradient from 6% to 35% solvent B. MS data were collected in positive ion, MS^E^ continuum, resolution mode with an m/z range of 50− 2000. Ion mobility was used to further resolve peptides in the gas phase. Peptides were fragmented by argon gas collision for data-independent acquisition. Peptic products were identified by fragmentation data using the ProteinLynx Global Server (PLGS version 3.0.3, Waters) with phosphorylation as an allowed variable modifier. Deuterium uptake was analyzed with DynamX version 3.0 (Waters). Peptides identified by PLGS were manually curated after establishing identifications thresholds in two of three unlabeled samples, minimum intensity of 4000, minimum sum of fragment intensities of 470, minimum score of 6.62, and 0.25 fragment per residue.^38^ Relative deuterium incorporation for each peptide and treatment was calculated by DynamX after manual inspection of spectra of each isotope envelope. Differences in deuterium incorporation between HDX treatments were assessed for representative time points approximating the actively exchanging regime of the time course. Statistical significance was determined using a two-tailed, unpaired t test. HDX-MS summary statistics are listed in Table S1 in accordance with HDX-MS community standards.^39^ The HDX-MS data and analysis files have been deposited to the ProteomeXchange Consortium via the PRIDE partner repository with the dataset identifier PXD034028.^40^ Exchange rate perturbations were mapped to Pyk2 structural models derived from AlphaFold Structure Database (AF-Q14289-F1)^29,41^ and ColabFold ver. 1.0.^30^

## Acknowledgements

We thank Joel Nott of the Iowa State University Protein Facility for mass spectrometry support. This research was funded by the National Science Foundation, Division of Molecular and Cellular Biosciences grant 1715411.

## Author Contributions

Conceptualization, E.S.U. and T.M.P.Z.; Methodology, E.S.U, T.M.P.Z., and A.W.; Investigation, T.M.P.Z., E.S.U, A.W., and K.G.R.; Writing – Original Draft, E.S.U., T.M.P.Z., A.W.; Writing – Review & Editing, E.S.U. T.M.P.Z., and A.W. Funding Acquisition, E.S.U.

