## Supporting Information for "Activation loop phosphorylation tunes conformational dynamics underlying Pyk2 tyrosine kinase activation"

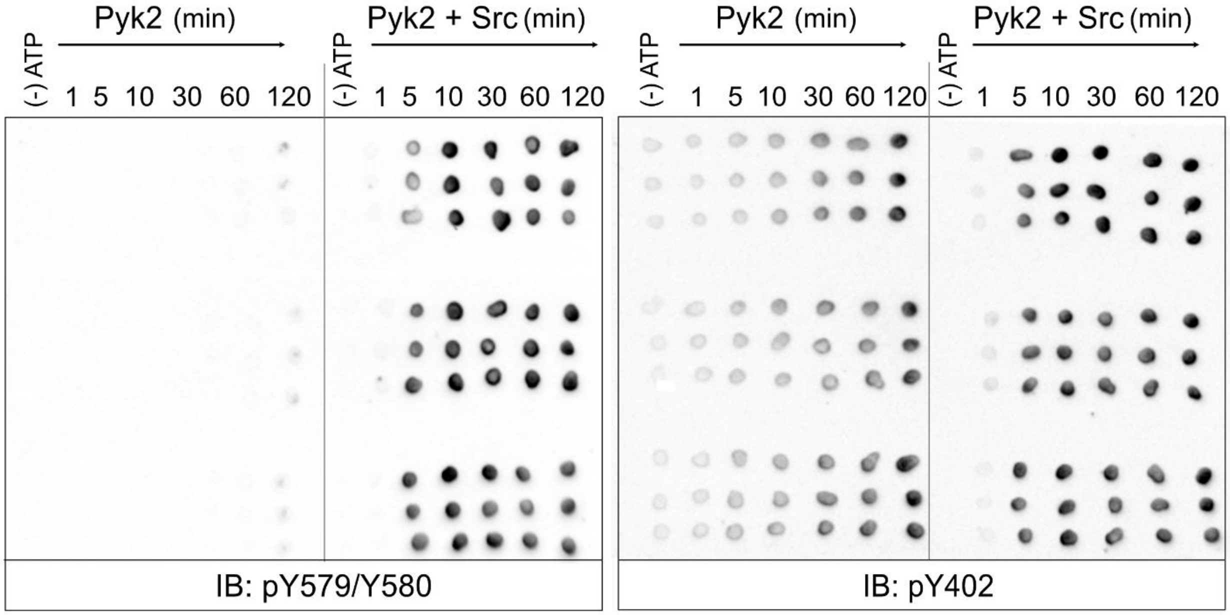


**Figure S1. Role of Src kinase in Pyk2 phosphorylation. Related to Figure 1.** Autophosphorylation (phospho-Y402) or Src-mediated activation loop phosphorylation (phospho-Y579/Y580) of Pyk2 FERM–kinase was initiated by addition of 4 mM ATP. Reaction time points were sampled over 120 min at 25°C. Time points were quenched with 100 mM EDTA and probed by immunoblotting with anti-phospho-PTK2B (pY402) or anti-phospho-PYK2 (pY579/Y580). Each row of dots represents one assay replicate time course.

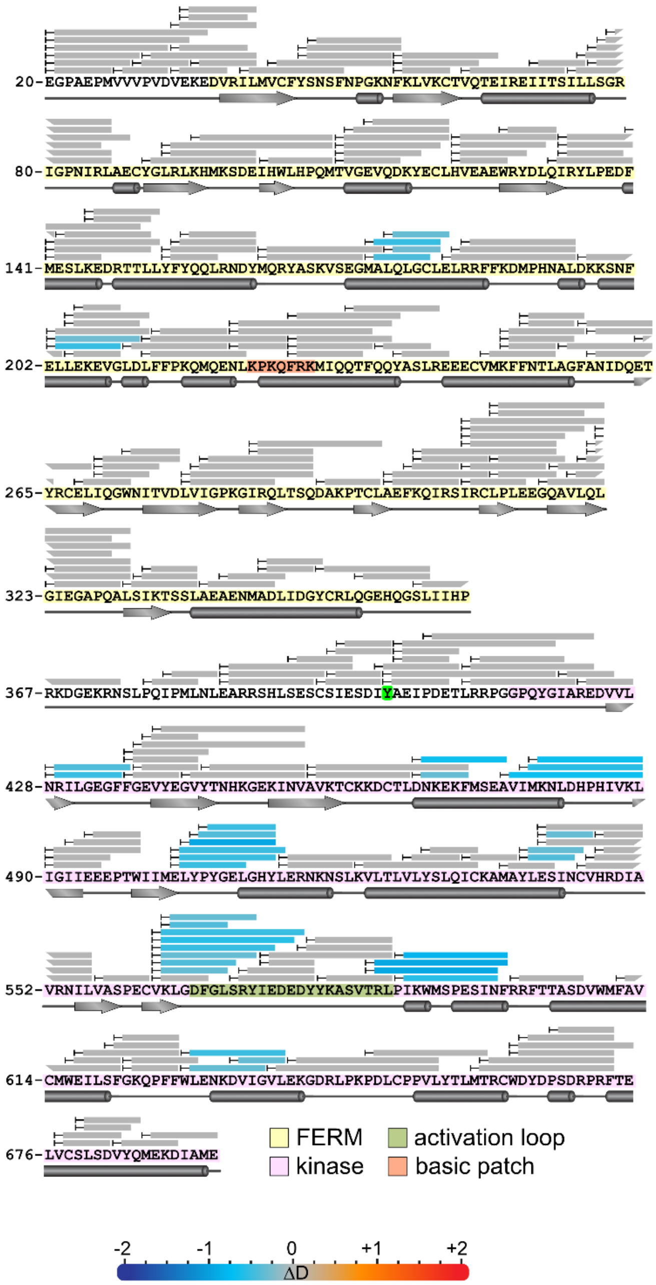


**Figure S2. Comparison of HDX-MS exchange kinetics of AMPPNP-bound Pyk2 FERM–kinase relative to unbound, Related to Figure 2.** Peptide coverage is represented as bars mapped to primary sequence. Exchange rate perturbations are reported as the average difference in deuterium incorporation (ΔD) at time points approximating the midpoint of exchange. Peptides exhibiting significant exchange rate perturbations are color-coded according to the scale bar (bottom). AMPPNP-induced decreases in H/D exchange are reported in blue shades, while increases in exchange are shown in reds. Significance was assessed with a two-tailed, unpaired Student's t-test ($p < 0.005$). Peptides exhibiting negligible differences are colored gray.

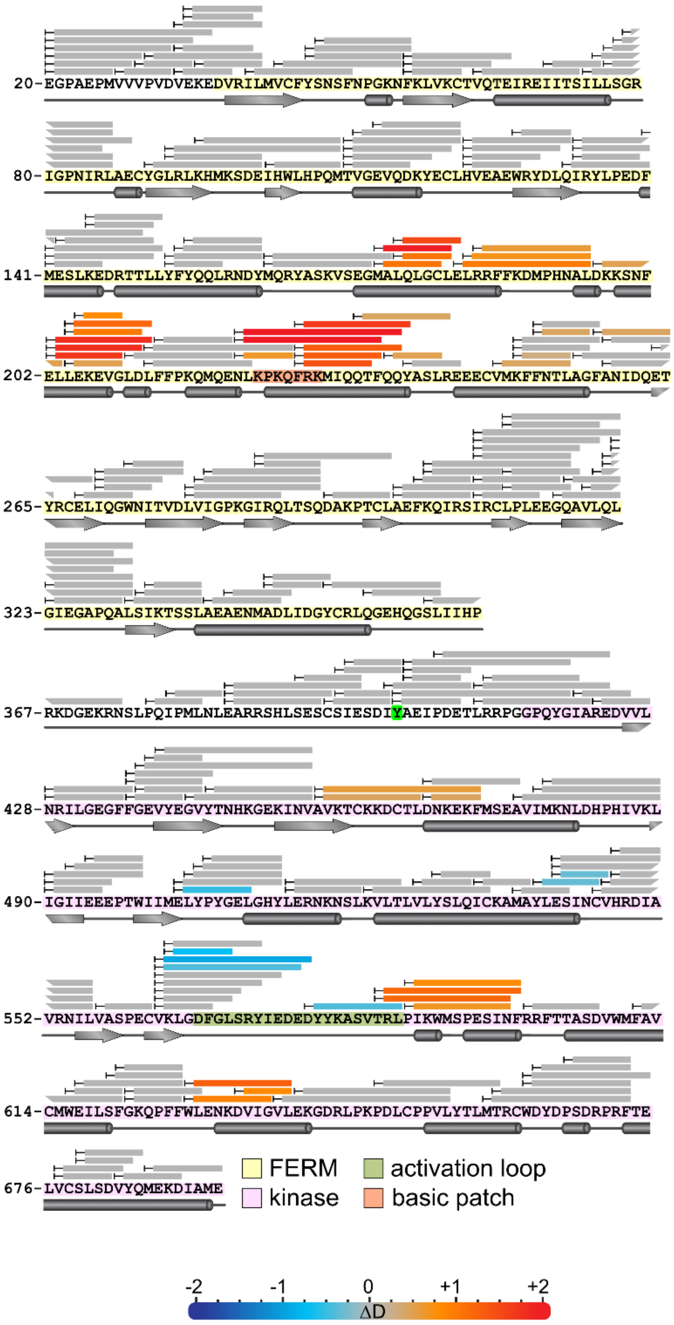


**Figure S3. Comparison of HDX-MS exchange kinetics of fully phosphorylated, ATP-bound Pyk2 FERM–kinase relative to AMPPNP-bound, Related to Figure 3.** Peptide coverage is represented as bars mapped to primary sequence. Exchange rate perturbations are reported as the average difference in deuterium incorporation (ΔD) at time points approximating the midpoint of exchange. Peptides exhibiting significant exchange rate perturbations are color-coded according to the scale bar (bottom). Phosphorylation-induced decreases in H/D exchange are reported in blue shades, while increases are shown in reds. Significance was assessed with a two-tailed, unpaired Student's t-test ($p < 0.005$). Peptides exhibiting negligible differences are colored gray.

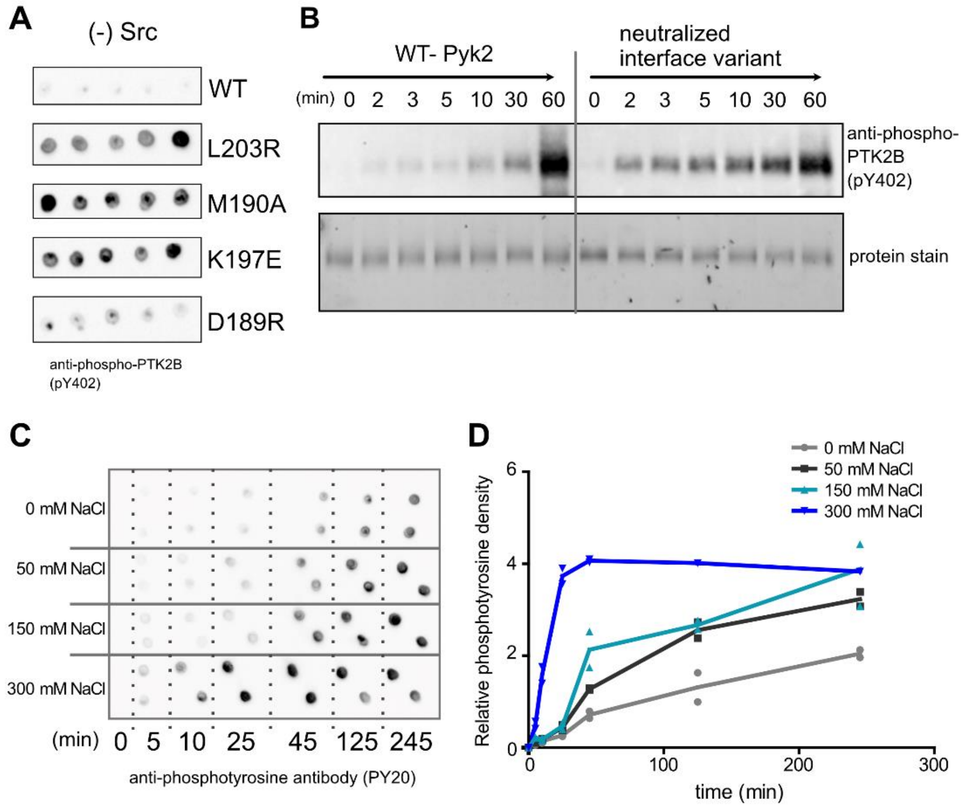


**Figure S4. Disruption of the Pyk2 FERM–kinase interface impacts Pyk2 kinase activity, Related to Figure 4.** (A) Autophosphorylation (5 min time point) of purified Pyk2 FERM–kinase constructs (WT vs FERM—kinase interface variants) was measured by dot blotting using anti-phospho-PTK2B (pY402). Each dot represents one reaction replicate. (B) Autophosphorylation activity time course of WT vs. neutralized interface variant (D189S, K197S, E207S, R600S, R601S, E639S) Pyk2 FERM–kinase constructs (0.5 µM) was initiated by adding ATP at 25°C and probed by immunoblotting with anti-phospho-PTK2B (pY402). Protein loading was imaged by 2,2,2-trichloroethanol staining.[1] (C) Autophosphorylation activity of WT Pyk2 FERM–kinase (1 µM) increases with increasing buffer ionic strength (0 – 300 mM NaCl). The kinase reaction was initiated by adding 4 mM ATP and quenched with 36 mM EDTA. Autophosphorylation was detected by dot immunoblotting using a pan-specific anti-phosphotyrosine primary antibody (PY20) and (D) quantified by densitometry.

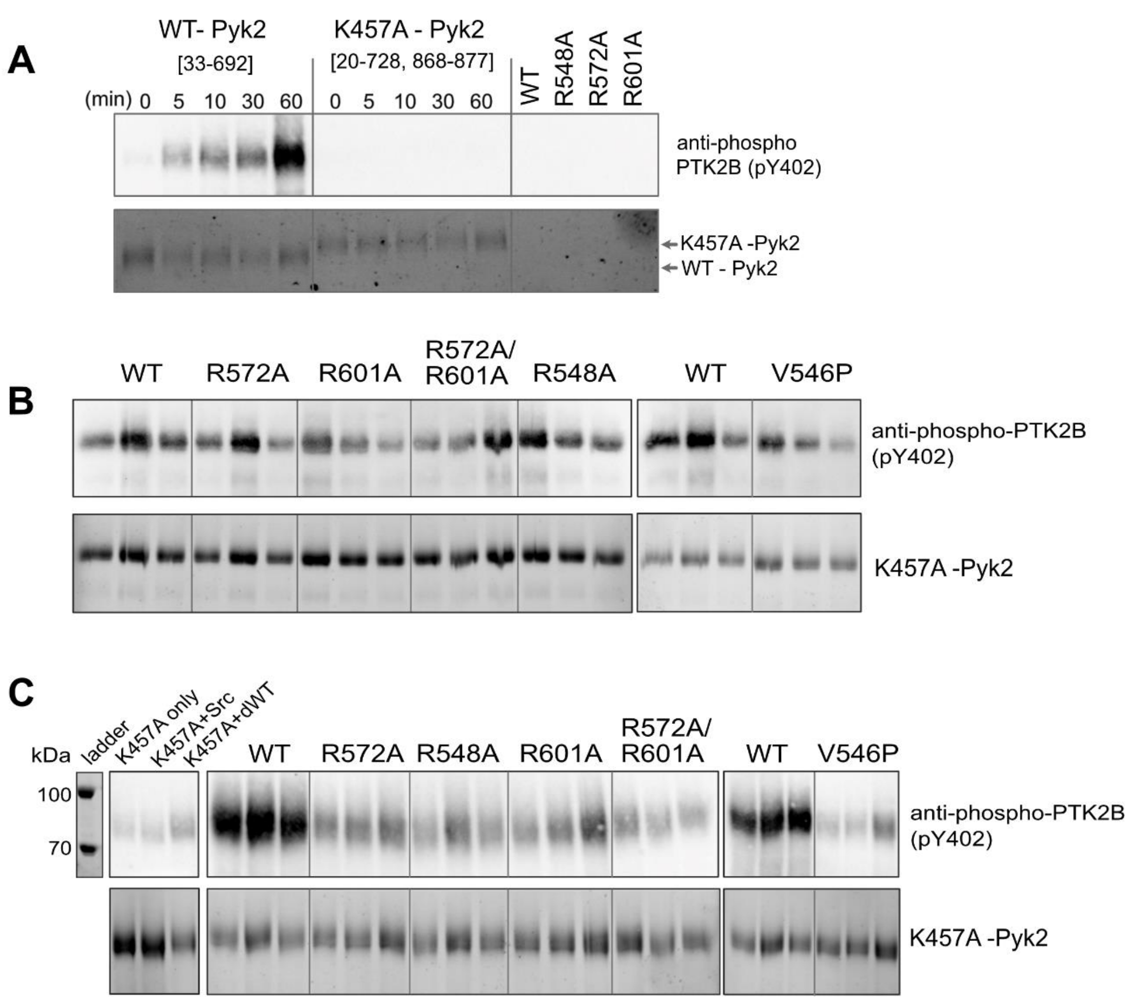


**Figure S5. Basic pocket and β6/β9 beta sheet are necessary for phosphorylation-induced high-activity conformation of Pyk2, Related to Figure 5.** (A) The kinase dead Pyk2 construct (K457A, residues 20-729/868-877) exhibits no detectable autophosphorylation activity as compared to WT Pyk2 FERM–kinase (1 µM) over a 60 min time course. Reactions were initiated with the addition of 4 mM ATP and quenched at various time points with 100 mM EDTA. Autophosphorylation of arginine variants at catalytic concentrations (0.02 µM) contribute negligible signal at the 60 min time point. (B-C) *In vitro* phosphorylation of K457A Pyk2 (1 µM) by catalytic amounts of unphosphorylated (B) or phosphorylated (C) Pyk2 FERM–kinase variants (0.02 µM) was carried out for 45 min and quenched with EDTA. Control experiments include K457A-Pyk2 alone (K457A only), K457A-Pyk2 with 0.006 µM Src (K457A+Src), and K457A-Pyk2 with 0.02 µM unphosphorylated WT Pyk2 (dWT). For (A-C), phosphorylation of K457A Pyk2 residue Y402 was measured by Western blotting using anti-phospho-PTK2B (pY402) primary antibody. Protein loading (bottom rows) was imaged by 2,2,2-trichloroethanol staining.[1]

**Table S1.** HDX-MS Data Summary Table. HDX analysis was performed by automated computational processing using DynamX software (Waters) and manual curation of each peptide, state, and time point. A data analysis summary is presented in the standardized format recommended by HDX community guidelines.[2]

| Data Set | Pyk2 FERM–kinase | Pyk2 FERM–kinase | Pyk2 FERM–kinase |
|---|---|---|---|
| | unbound | +AMPPNP | +ATP + Src |
| HDX reaction details | 50 mM HEPES pD 7.40, 100 mM NaCl, 5 mM DTT, 5 mM $MgCl_2$, 90% $D_2O$, 21.8 °C | 50 mM HEPES pD 7.40, 100 mM NaCl, 5 mM DTT, 5 mM $MgCl_2$, 90% $D_2O$, 21.8 °C | 50 mM HEPES pD 7.40, 100 mM NaCl, 5 mM DTT, 5 mM $MgCl_2$, 90% $D_2O$, 21.8 °C |
| HDX time course (min) | 0.2, 1, 2.5, 11, 122 | 0.2, 1, 2.5, 11, 122 | 0.2, 1, 2.5, 11, 122 |
| HDX control samples | undeuterated controls, n=3 | | |
| Back-exchange | ND | | |
| # of Peptides | 264 | 264 | 258 |
| Sequence coverage | 98.10% | 98.10% | 98.10% |
| Average peptide length / Redundancy | 8.5/3.9 | 8.5/3.8 | 8.6/3.9 |
| Replicates | 3 (technical) | 3 (technical) | 3 (technical) |
| Repeatability (Average std. dev. of D uptake) | 0.082 | 0.083 | 0.086 |
| Significance testing | two-tailed, unpaired t test, p<0.005 at time point(s) approximating the middle range of exchange | | |

**Table S2.** List of oligonucleotide primers used in this study. Details described the plasmid identity and the mutation to the cDNA sequence.

| Name | Sequence (5'-3') | Details |
|---|---|---|
| TP020 | GTAGCCGTAGCAACCTGTAAAAAGGAC | pAAW001/ K457A |
| TP021 | GTCCTTTTTACAGGTTGCTACGGCTAC | pAAW001/ K457A |
| TP030 | GATAAAAAGAGCAACTTCGAAAGGCTTGAGAAAGAGGTTGGG | pNAM004/L203R |
| TP031 | CCCAACCTCTTTCTCAAGCCTTTCGAAGTTGCTCTTTTTATC | pNAM004/L203R |
| TP032 | GCGTCGCTTCTTTAAGCGCATGCCGCACAACGCTC | pNAM004/D189R |
| TP033 | GAGCGTTGTGCGGCATGCGCTTAAAGAAGCGACGC | pNAM004/D189R |
| TP034 | CGCACAACGCTCTGGATGAAAAGAGCAACTTCGAAC | pNAM004/K197E |
| TP035 | GTTCGAAGTTGCTCTTTTCATCCAGAGCGTTGTGCG | pNAM004/K197E |
| TP052 | GCTTCTTTAAGGACGCGCCGCACAACGCTC | pNAM004/M190A |
| TP053 | GAGCGTTGTGCGGCGCGTCCTTAAAGAAGC | pNAM004/M190A |
| TP054 | GAATCTATCAACTGTGTGCATGCTGACATTGCTGTGCGTAAT | pNAM004/R548A |
| TP055 | ATTACGCACAGCAATGTCAGCATGCACACAGTTGATAGATTC | pNAM004/R548A |
| TP058 | GGGAGACTTTGGATTAAGTGCTTACATTGAGGATGAGGAC | pNAM004/R572A |
| TP059 | GTCCTCATCCTCAATGTAAGCACTTAATCCAAAGTCTCCC | pNAM004/R572A |
| TP082 | CTTGAATCTATCAACTGTCCCCATCGTGACATTGCTGTG | pNAM004/V546P |
| TP083 | CACAGCAATGTCACGATGGGGACAGTTGATAGATTCAAG | pNAM004/V546P |
| TP084 | GAGTCTATCAATTTTCGCGCTTTTACCACGGCTTCGGATGTC | pNAM004/R601A |
| TP085 | GACATCCGAAGCCGTGGTAAAAGCGCGAAAATTGATAGACTC | pNAM004/R601A |
| TP116 | CAATTTTAGCAGTTTTACCACGGCTTCG | pNAM004- R600S/R601S |
| TP118 | ATAGACTCCGGGGACATCCACTTAATTGGC | pNAM004- R600S/R601S |
| TP121 | GCTCTGGATTCAAAGAGCAACTTCGAACTGCTTGAG | pNAM004/K197S |
| TP122 | GTTGTGCGGCATGGACTTAAAGAAGCGAC | pNAM004/D189S |
| TP119 | GTTATCGGCGTACTGTCAAAGGGCGATCGTCTTCCG | PNAM004/E639S |
| TP120 | CGGAAGACGATCGCCCTTTGACAGTACGCCGATAAC | PNAM004/E639S |
| TP125 | GAACTGCTTGAGAAATCGGTTGGGTTGGACTTGTTTTTCCCC | PNAM004/E207S |
| TP126 | GGGGAAAAACAAGTCCAACCCAACCGATTTCTCAAGCAGTTC | PNAM004/E207S |
| TP123 | GCGTCGCTTCTTTAAGAGCATGCCGCACAACGCTC | PNAM004/D189S |
| TP124 | GAGCGTTGTGCGGCATGCTCTTAAAGAAGCGACGC | PNAM004/D189S |
